# microntology: a lightweight, data-driven controlled vocabulary to describe Earth’s microbial habitats

**DOI:** 10.64898/2026.01.12.698811

**Authors:** Anthony Fullam, Vishnu Prasoodanan PK, Michael Kuhn, Peer Bork, Thomas SB Schmidt

## Abstract

**Summary:** We introduce *microntology* v1.0, a pragmatic controlled vocabulary of 148 terms to describe microbial habitats and lifestyles, and provide manually curated *microntology* annotations for >300k metagenomic samples from public repositories.

**Data Availability:** *microntology* controlled vocabulary terms and term hierarchies ^1^, and curated annotations for 305,626 metagenomic samples ^2^ are available via Zenodo and as supplemental tables S1 and S2. User feedback, suggestions and bug reports are welcome under https://github.com/grp-schmidt/microntology/issues.

## Main Text

Data on microbial life continues to accrue in public repositories: at the time of writing, nearly 6 million amplicon^1^ sequencing and 1.2 million metagenomic^2^ experiments on microbial communities are freely accessible via the International Nucleotide Sequence Database Collaboration (INSDC, ^3^), with rapid growth also in other ‘omic data types ^4,5^. This wealth of information enables studies of microbial ecology and evolution at unprecedented *depth* (i.e., with deep readouts on individual samples or habitats) and *breadth* (across many samples, often across different habitats), facilitated by dedicated resources that integrate microbial data across Earth’s biomes, such as catalogues of genes ^6^, gene families ^7^, metabolic pathways ^8^, metagenome-assembled genomes (MAGs, ^9–12^) or taxonomic profiles ^13,14^.

A key challenge in navigating repositories and identifying datasets relevant to specific research questions is the varying availability of consistent contextual data, which is why INSDC databases require submissions to conform with MIxS standards by providing Minimum Information about Genome/Metagenome Sequences (MIGS/MIMS, ^15^), MAGs (MIMAGS, ^16^) or MARKer gene Sequences (MIMARKS, ^17^). These include descriptions of the broad-scale and local environmental context (formerly defined as annotation fields ‘environment (biome)’ and ‘environment (feature)’), as well as the sampled environmental medium (formerly ‘environment (material)’), with a recommendation to use terms from corresponding classes in the Environment Ontology (EnvO, ^18,19^). However, annotations are often incomplete, uninformative (using generic terms) or indeed erroneous ^20^ and require extensive curation and harmonization, provided via dedicated databases such as curatedMetagenomicData ^21^ or Metalog ^20^. Moreover, EnvO ^18,19^ and other available controlled vocabularies to describe microbial habitats, such as Omnicrobe ^22^ or the GOLD Ecosystem Classification ^23^ used for IMG/M resources ^24^, are designed towards maximum resolution, containing thousands of classes that can be difficult to navigate and summarise into broader categories of related terms.

Here, we introduce *microntology*, a lightweight controlled vocabulary to describe microbial habitats and lifestyles. The *microntology* follows five core design principles: it is (i) *data-driven*, encompassing terms to describe frequent sample types in existing data; (ii) *pragmatic and query-oriented*, defining terms at intermediate resolutions to facilitate user access and browsing; (iii) *shallow*, with only few hierarchical levels of classes; (iv) *cross-linked* between related terms from different hierarchical paths (e.g., a lake [MICRONT:02010100] would also be considered a lentic water body [MICRONT:02060100]); and (v) designed for *parsimonious multi-tagging* where individual samples are described by multiple independent terms, rather than a ‘best match’ highly detailed single term (see examples below).

In v1.0, the *microntology* encompasses a controlled vocabulary of 148 terms that fall into five broad habitat categories (terrestrial, aquatic, aerial, host-associated and anthropogenic environments) and additional categories that describe broad physicochemical properties (sample salinity, temperature, pH and oxygen level which are often associated with microbial community structure in different habitats), morphology (biofilms), human host properties to further differentiate within this dominant sample group (age group, birth term, place of residence, non-industrialized lifestyles), animal host captivity status, and miscellaneous informative features such as (chemical) contamination or ancient (paleontological) sample provenance. Terms are moreover linked to matching classes in EnvO ^18,19^, UBERON ^25^ and other existing ontologies, as well as to NCBI Taxonomy IDs ^26^ where applicable. An overview of *microntology* terms (and their frequency among currently available metagenomes, see below) is shown in Figure 1; a full table including resolved hierarchies, cross-linked terms and definitions is available as Supplementary Table S1 and in ^1^.

**Figure 1.**
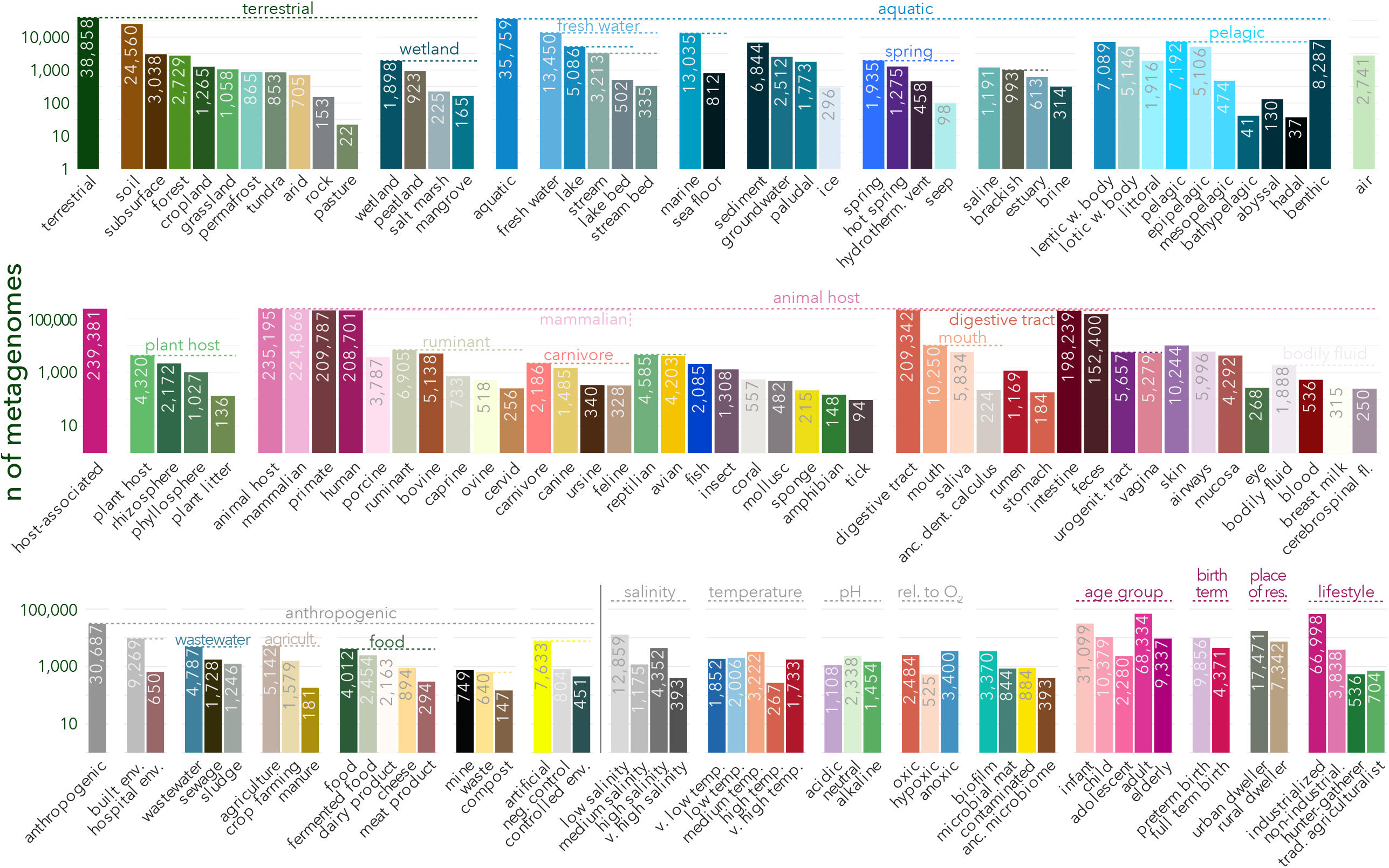
Overview of curated microntology annotations for >300k publicly available metagenomes. For each *microntology* term (categorical x axis), the number of annotated metagenomic samples (y axis) is shown. Categories are organized by sample group. Note that y axis scales differ between rows for improved visibility. As annotation follow hierarchical categories and individual samples are annotated with multiple tags (see main text), total counts do not sum to the total number of 305,626 annotated metagenomes.

Major habitat categories are further resolved along parallel yet cross-linked paths. Terrestrial habitats (MICRONT:01000000) can be described based on sample type (e.g., soil, rock, subsurface) and environmental context (forest, grassland, tundra, etc). Following *microntology’s* underlying philosophy of using multiple descriptive tags, a forest soil sample would be annotated using the terms ‘soil’ (MICRONT:01010000) and ‘forest’ (MICRONT:01020400), rather than a specific term ‘forest soil’ as a subclass of one or the other. Aquatic habitats (MICRONT:02000000) are further categorized based on general salinity ecosystem (fresh water, brackish, marine, non-marine saline, brine), water body type (lentic or lotic), broad water layer (littoral, pelagic, benthic), and dedicated categories describing springs (including hydrothermal vents), ice, and groundwater samples. Aquatic sediments are considered as a distinct subcategory, crosslinked to benthic features (lake and stream beds, seafloor). Descriptions of habitats at terrestrial-aquatic interfaces are resolved via both crosslinking of terms and multiple tagging: ‘groundwater’ (MICRONT:02070100) is considered an aquatic habitat, but automatically crosstagged as terrestrial and ‘subsurface’ (MICRONT:01030000); ‘wetland’s (MICRONT:01020700) are categorized as terrestrial, but crosstagged as ‘lentic’ (MICRONT:02060100) and ‘paludal’ (MICRONT:02090000) water bodies; soils from intertidal zones or ocean coasts would in practice be co-tagged as ‘littoral’ (MICRONT:02080100) and ‘marine’ (MICRONT:02020000).

Host-associated samples (MICRONT:03000000) are broadly split into ‘plant host’ (MICRONT:03010000) and ‘animal host’ (MICRONT:03020000) groups. The latter is further resolved based on broad host taxonomy (mammalian, avian, sponge hosts, etc), body site (digestive tract, urogenital tract, skin, airways, etc) and sample type (bodily fluids or mucosa). Feces (MICRONT:03030141), saliva (MICRONT:03030111) and ancient dental calculus (MICRONT:03030112) are sample types that are treated as special cases, as they are unambiguously intestinal (MICRONT:03030140) and oral samples (MICRONT:03030110), respectively. Following *microntology’s* multi-tag approach, a dog fecal sample would be annotated as ‘canine’ (MICRONT:03020131) and ‘feces’ (MICRONT:03030141), whereas caecal contents of chicken would be considered as ‘avian’ (MICRONT:03020210) ‘intestine’ (MICRONT:03030140). For ‘human’ (MICRONT:03020111) samples, by far the most data-rich category in practice, additional information can be added by tagging ‘age group’ (from infant to elderly), ‘birth term’ (full or preterm birth), ‘place of residence’ (urban or rural) and ‘host lifestyle’ (e.g., non-industrialized).

The ‘anthropogenic’ (MICRONT:04000000) category includes engineered or directly human-impacted habitats, such as the ‘built environment’ (MICRONT:04020000), ‘agriculture’-associated environments (MICRONT:04040000), ‘wastewater’ (MICRONT:04030000) and ‘food’ (MICRONT:04060000). The *microntology* includes a special ‘artificial sample’ category (MICRONT:04010000) to flag mock communities or negative controls, but also to describe experimentally derived samples from raw environmental starting materials, e.g. using controlled environments, cell sorting or enrichment cultures. For example, microcosm experiments from lake sediments would be annotated as ‘lake bed’ (MICRONT:02010101), but qualified with a ‘controlled environment’ (MICRONT:04010300) tag.

We curated *microntology* annotations for 305,626 publicly available shotgun metagenomic samples from 1,688 studies (Figure 1; per-sample data available under ^2^). We initially included all metagenomes (*source: METAGENOMIC* and *strategy: WGS*) from projects containing ≥20 samples with ≥1M reads and ≥1Gpb sequencing depth, generated on Illumina machines or compatible platforms (DNBseq, BGIseq), that were available on the European Nucleotide Archive (ENA, ^27^) on 31st October 2024. From this initial list, datasets were excluded upon manual inspection if they were (i) misannotated amplicon or genomic sequencing data; (ii) artificial samples such as synthetic metagenomes or mock communities; (iii) obtained from *in vivo* models in a laboratory context, e.g. mouse experiments; (iv) enriched for viral particles or specific prokaryotic community members. Additional datasets were included on a per-study basis, in particular from IMG/M ^24^ and SPIRE ^11^. For 1,501 studies (representing 95% of samples), ENA project IDs were matched to publications following a previously described approach ^11,20^. *microntology* terms were annotated in multiple steps. First, for a subset of 129,904 samples, available manually curated information was parsed from Metalog ^20^ and matched to *microntology* tags. Next, we mapped tags to entries from ENA contextual data fields (such as environmental context, scientific name and host tax ID). Finally, for a subset of 163 studies and an additional 22,862 samples, *microntology* tags were manually added based on information in associated publications or additional (not automatically parsed) ENA contextual data fields.

In our survey (Figure 1; see Figure S1 for an alternative view), publicly available metagenomic data remains strongly biased towards animal-associated habitats (235k samples, or 77% of total), and in particular to human (209k, 68.5%) and intestinal (198k, 65%) samples. However, the representation of other body sites (such as mouth, skin, airways and urogenital tract), non-human hosts (in particular bovine, porcine and avian), but also of soils (25k samples), fresh water (13.5k), ocean (13k), the built environment (9k) and food-associated habitats (4k) has increased significantly compared to recent surveys (e.g., ^11^). We note that for additional descriptive categories, such as salinity, temperature or human age group or place of residence, annotation recall is necessarily limited, as the corresponding information is often not available in public repositories, nor can it be extracted from publications. In such instances, we annotated conservatively, i.e. assigning only tags unambiguously supported by publicly available data, without extrapolation.

*microntology* annotations have previously been used to facilitate data exploration e.g. in SPIRE v1 ^11^ and the Global Microbial SmORFs Catalogue ^28^, to benchmark global-scale microbial habitat unsupervised taxonomic clusterings and modeling ^29^, and to categorize the main reservoirs of discoverable microbial diversity in existing data ^30^. The resource is continuously developed and iteratively refined, following the basic principles laid out above, i.e. driven by data availability and user requirements in practice, relying on multiple parsimonious tags to describe microbial habitats at an intermediate and pragmatic resolution. The *microntology* is intended to provide a lightweight complement to established highly detailed ontologies such as EnvO ^19^ or the GOLD Ecosystem Classification ^31^ while retaining interoperable links, yet to provide broader coverage than project-specific vocabularies such as the Earth Microbiome Project ^32^ or Microflora Danica ontologies ^33^. *microntology* annotations can be browsed as a rough classification of datasets towards explorative analyses, although detailed contextual data for finescale analyses needs to be retrieved from dedicated resources, such as curatedMetagenomicData ^21^ or Metalog ^20^. In summary, *microntology* annotations may facilitate studies of microbial ecology and evolution at large scales, integrated across Earth’s microbial habitats and lifestyles.

## Acknowledgements

The authors thank Luis Pedro Coelho (Queensland University of Technology) for helpful discussions and feedback. This research was conducted with the financial support of Research Ireland under Grant Number 12/RC/2273-P2 (V.P.P.K and T.S.B.S).

---

**Figure S1.**
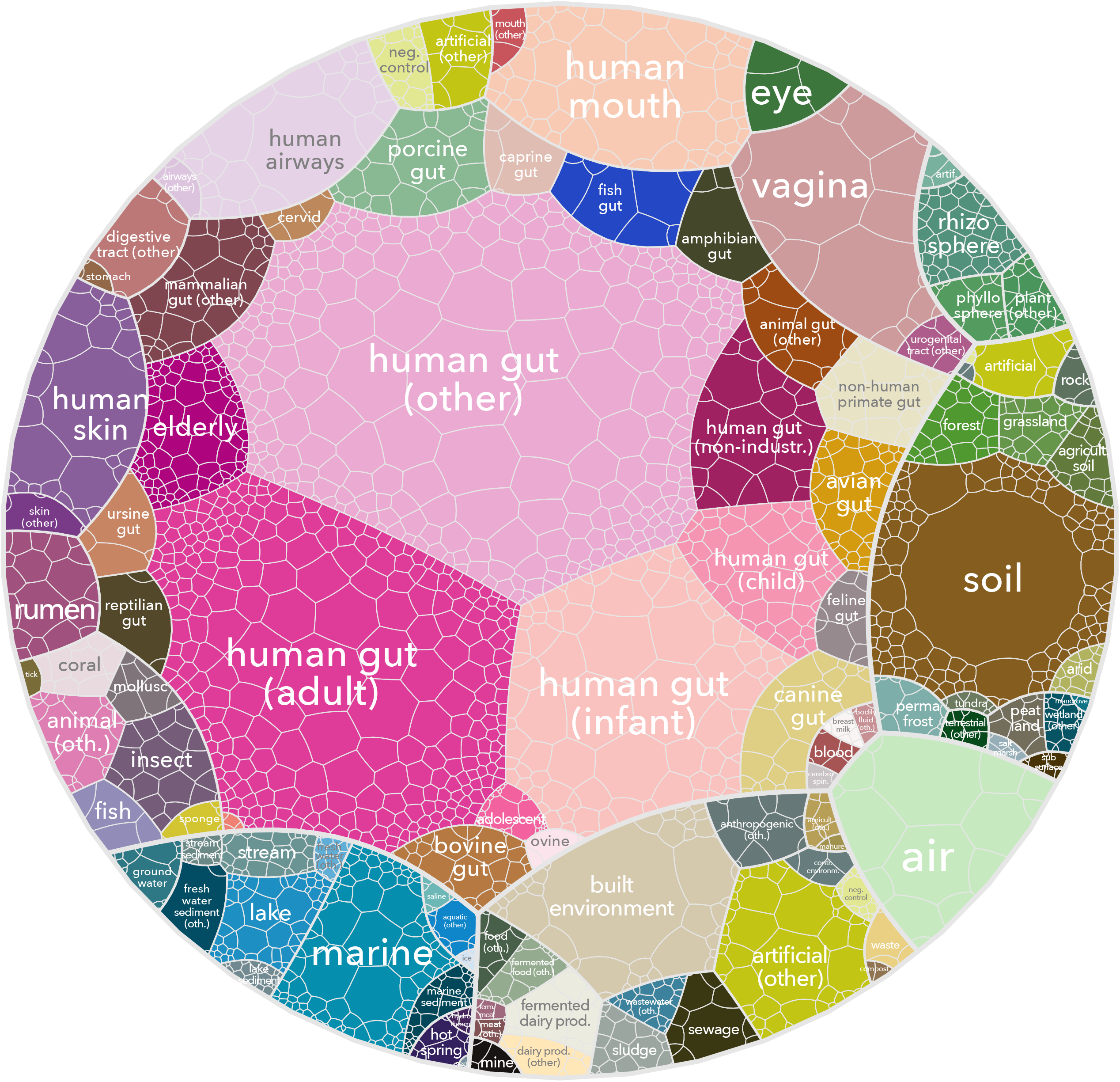
A habitat-resolved view of available metagenomic data. *microntology* annotations were further summarised to label each sample under a most descriptive category. In the treemap plot, each cell corresponds to an individual study (out of 1,688 total studies); cell size corresponds to the number of samples in a study in the focal habitat category.

## Footnotes

1 ncbi.nlm.nih.gov/sra, accessed Dec 16th 2025; query *(“metagenomic”[Source]) AND “amplicon”[Strategy]*

2 query *(“metagenomic”[Source]) AND “wgs”[Strategy]*

## Notes

https://zenodo.org/records/18162129

https://zenodo.org/records/18164252

### Competing Interest Statement

The authors have declared no competing interest.

